# Thermodynamic Measures of Human Brain Development from Fetal Stage to Adulthood

**DOI:** 10.1101/547364

**Authors:** Edward A. Rietman, Sophie Taylor, Hava T. Siegelmann, Marco Cavaglia, Jack A. Tuszynski

**Affiliations:** BINDS Lab, School of Computer Science, University of Massachusetts Amherst, 140 Governors Drive, Amherst, MA, 01003-9264, United States; Department of Physics, University of Alberta, 4-181 CCIS, Edmonton, AB, T6G 2E1, Canada; ACTISMED, srl, Torino, Italy; DIMEAS, Politecnico di Torino, Corso Duca degli Abruzzi 24, Torino, 10129, Italy; Department of Oncology, University of Alberta, Cross Cancer Institute, 11560 University Avenue, Edmonton, AB, T6G 1Z2, Canada

**Author notes:** Corresponding author: Jack A. Tuszynski, 3-336, Cross Cancer Institute, 11560 University Avenue, Edmonton, AB, T6G 1Z2, Canada.

**Keywords:** brain development, Gibbs free energy, thermodynamics, transcriptome, protein-protein interaction

## Abstract

This paper analyzes the data obtained from tissue samples of the human brains containing protein expression values. The data have been processed for their thermodynamic measure in terms of the Gibbs free energy of the corresponding protein-protein interaction networks. We have investigated the functional dependence of the Gibbs free energies on age and found consistent trends for most of the 16 main brain areas. The peak of the Gibbs energy values is found at birth with a trend toward plateauing at the age of maturity. We have also compared the data for males and females and uncovered functional differences for some of the brain regions.

**Significance Statement:** In this paper we briefly outline the theoretical basis for a novel analysis of brain development in terms of a thermodynamic measure (Gibbs free energy) for the corresponding protein-protein interaction networks. We analyzed the overall developmental patterns for Gibbs free energy as a function of age across all brain regions. Of particular note was the significant upward trend in the fetal stages, which is generally followed by a sharp dip at birth and a plateau at maturity. We then compared the trends for female and male samples. A crossover pattern was observed for most of the brain regions, where the Gibbs free energy of the male samples were lower than the female samples at prenatal and neonatal ages, but higher at ages 8-40.

## Introduction

In this paper, we introduce a possible thermodynamic measure that could correlate with brain development from the fetal stage to old age as measured by protein expression data for all major brain regions. The thermodynamic measure we have used in this study is the Gibbs free energy of the protein-protein interaction network. This approach is motivated by previous applications of the Gibbs energy of such networks as a measure of biological state evolution. In particular, we earlier showed that the Gibbs free energy calculated for the development of *C. elegans* is correlated with the developmental timeline of this species (1). The theoretical underpinnings for understanding the thermodynamics and energetics of *C. elegans* development or brain development, started by investigating the molecular biology of human diseases from a systems biology perspective. These studies were carried out over a several-year period (2–6). From a thermodynamic perspective, the transcriptome and other omic (e.g., proteomic, genomic, metabolomic, etc.) measures can represent the energetic state of a living cell or indeed an organism as a whole. There is a chemical potential associated with the interacting molecules in a cell, and the chemical potential of all the proteins that interact with each other can be viewed to represent a rugged landscape, not dissimilar to Waddington’s epigenetic landscape (7, 8). From a biological perspective, we know that 74% of the human transcriptome is over-expressed in the brain compared to other organs, meaning that 14,518 out of 19,614 of all human proteins; and hence some 14,600 of the corresponding genes show an elevated expression in the brain compared to other tissue and organ types (9, 10).

A previous study based on gene ontology analysis and antibody-based tissue microarray analysis of the corresponding proteins, found most brain-enriched protein coding genes to be found in astrocytes, oligodendrocytes or in neurons with molecular properties linked to synaptic transmission and brain development (11). Moreover, recent findings show that approximately 86% of protein-coding genes are differentially regulated at the whole transcript or exon level across regions and/or time and most spatio-temporal differences occur before birth, followed by an increase in the similarity among regional transcriptomes during postnatal lifespan. These findings demonstrate that genes are organized into functionally distinct co-expression networks, and sex differences are present in gene expression and exon usage (12). In recent years biophysical studies have been providing thermodynamic interpretation of biological processes adding the necessary conceptual framework which helps to navigate the inherent complexities and better understand biology without including superfluous or misleading information (13). The method we propose here demonstrates that the thermodynamic spatio-temporal profile of the human brain transcriptome, when compared with biological data from neurobiology literature, correlates from the DNA level all the way to the organ level. As will be shown below, the most important findings of our work are that the human brain’s transcriptome has the lowest (most negative) Gibbs free energy values pre-birth, followed by a dramatic climb to the maximum Gibbs free energy at birth, which mirrors not only the growth of the organ itself but also the development of its complex internal architecture. This rapid ascent to a maximum value is then followed a gradual drop in the Gibbs free energy to a local minimum around the age of sexual maturity. After that, another change in the trend occurs with a slight increase in the Gibbs energy continuing into the old age. These pronounced trends, which are consistent across the various areas of the brain, separated by rather sharp transitions, in our opinion related to important biological and physiological processes such as structure formation, building of neuronal connections and also structural and functional deterioration due to aging. Interestingly, there are also documented thermodynamic transitions taking place in the human brain such as the sudden temperature change at birth, which corresponds to a change of conditions from a thermodynamically closed system to an open one. The passage to old age is also known to correspond to volumetric changes of the brain as well as pathological protein aggregation such as seen in the formation of amyloid plaques (14).

## Data Sources and Methods

The method we propose uses mRNA transcriptome data or RNA-Seq data as a surrogate for protein concentration. This assumption is largely valid. Kim et al. (15) and Wilhelm et al. (16) have shown an 83% correlation between mass spectrometry proteomic information and transcriptomic information for multiple tissue types. Further, Guo et al. (17) found a Spearman correlation of 0.8 in comparing RNA-Seq and mRNA transcriptome from TCGA human cancer data (18). We believe this justifies the use of this methodological simplification, especially since we are interested in trends and not in the quantification of minute differences.

Given the set of transcriptome data, a representative of protein concentration, we then overlay that on the human protein-protein interaction network from BioGRID (19, 20). This means we assign to each protein on the network, its scaled (between 0 and 1), transcriptome value (or RNA-Seq value). From that we compute the Gibbs free energy of each protein-protein interaction using the relationship:

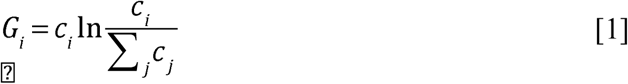

where *c*_*i*_ is the “concentration” of the protein *i*, normalized, or rescaled, to be between 0 and 1. The sum in the denominator is taken over all protein neighbors of *i,* and including *i.* Therefore, the denominator can be considered related to a degree-entropy of the underlying protein-protein interaction network. Carrying out this mathematical operation essentially transforms the “concentration” value assigned to each protein to a Gibbs free energy contribution, which can eventually be added up for the entire set of expressed proteins providing an overall thermodynamic measure for the state of the system at a given point in time. Thus, we replace the set of scalar values associated with a transcriptome by a scalar function – the Gibbs free energy. By summing over the whole protein-protein interaction network (PPI), we can compute the Gibbs free energy, which represents the energetic state of the biopsy tissue sample.

The transcriptome data provide quantitative information about the mRNA expression levels in a given sample. Expression values are log2 normalized. These expression levels, *e*_*i*_, act as a proxy for protein concentration. The concentrations values, *c*_*i*_, are further normalized into the interval [0,1] where a value of 1 suggests a high relative protein concentration, *e*_*max*_, while a value of 0 corresponds to a low relative protein concentration, *e*_*min*_, according to

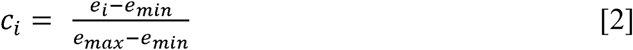

With these two sets in hand, we superimpose the protein concentration data on the PPI. A chemical potential is then assigned to each node using the relationship:

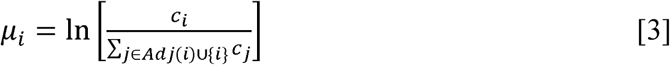

where the denominator is the sum of the concentrations of adjacent nodes. The term in the argument of the natural logarithm function will be equal to 1 if the protein is unconnected (or all adjacent nodes have 0-concentraction values). Generally, we can see that µ _*i*_ will be greater for higher values of *c*_*i*_ and lower concentrations in adjacent nodes. The magnitude of the potential is therefore greater for low concentration proteins with highly upregulated adjacent nodes.

We can then perform a simple calculation to obtain the total Gibbs free energy for the network

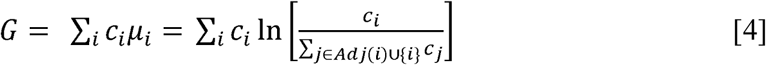

We performed these calculations for several transcriptome data sets, however the focus of this paper is specifically on the data set GSE25219 (12, 21) from the NCBI GEO database (22, 23), which contains 1,340 samples obtained from 57 healthy postpartum human brains. The data set contains quantitative information for 16 brain regions with samples including both sexes and multiple age groups ranging from prenatal to adult (12). A list of these brain regions and their acronyms used can be found in Table 1 (12). The PPI data were obtained from BioGRID (19, 20). The Gibbs energy was computed using Python 2.7. Biological interpretation of thermodynamic findings has been carried out by reviewing the relevant scientific literature.

**Table 1.** The list of brain regions used in the study of the human brain development (12).

| Abbreviation | Brain region |
| --- | --- |
| OFC | orbital prefrontal cortex |
| DFC | dorsolateral prefrontal cortex |
| VFC | ventrolateral prefrontal cortex |
| MFC | medial prefrontal cortex |
| M1C | primary motor (M1) cortex |
| S1C | primary somatosensory (S1) cortex |
| IPC | posterior inferior parietal cortex |
| A1C | primary auditory (A1) cortex |
| STC | posterior superior temporal cortex |
| ITC | inferior temporal cortex |
| V1C | primary visual (V1) cortex |
| HIP | hippocampus |
| AMY | amygdala |
| STR | striatum |
| MD | mediodorsal nucleus of thalamus |
| CBC | cerebellar cortex |

Before, we present the results of this analysis, we briefly outline the workflow involved in the data analysis. We downloaded the data from the following data sets from GEO using the GPL5175 platform (Affymetrix Human Exon 1.0 ST Array): GSE30272 (24), GSE18069 (25), and GSE25219 (12, 21), as well as BrainSpan (26, 27). Only GSE25219 was selected for further analysis. GSE25219 contained 57 subjects, 16 brain regions, LR hemispheres (for only 39 of the subjects), 31 males and 26 females. The samples obtained were from normal donors without a clinical brain pathology or signs of serious genomic abnormalities (12). We then combined soft matrix data with gene expression data files. IDs in the gene expression data were matched to gene names (several ID numbers can match to a single gene name). Where there were duplicate entries for a gene, the average of the expression values was taken. If there was no gene name that could be matched to an ID, the entry was deleted. The output file created contains a list of genes (rows) and samples (1,340 columns) with the averaged log2 expression values for each entry. We then calculated Gibbs free energy values associated with each sample and plotted the results (see below) by binning data into age groups. The age ranges were determined based on natural breaks in the data set, developmental stage, and considerations based on the amount of data in a given set (for example, a bin might be increased if there was insufficient data within the bin). The difference in binning ranges between plotting sets (i.e. sex, protein, overall, hemisphere) is due to this third factor. Error bars are based on standard error calculations. Some of the fetal regions used a different labelling system. In our analysis, any data that were labelled using a label that could not be directly matched to one of the 16 brain regions mapped were discarded. Fortunately, this constituted a very small fraction. All of the final plots were produced using the R software, along with the accompanying statistical data.

## Results

In this section we provide an overall transcriptome time dependence of the corresponding Gibbs free energy and a discussion of the physiological interpretation when available. Figure 1 shows the overall patterns in the 16 regions of the human brain from prenatal stages to old age. In Figure 1, a simple average has been taken for all samples of a particular brain region for a given age range (with suppressed hemisphere and sex information). We can observe a consistent upward trend in the fetal period across all brain regions. The three regions that do not appear to increase between the first and second time periods are V1C, S1C, and M1C, and the upper outlier in the second bin is MD. The data are in accordance with the embryologic development time of V1C, S1C, and M1C not yet present (28). The high values of the Gibbs free energy for MD, on the contrary, are related to its earlier and complex development, starting immediately after the formation of the neural tube (29, 30). The data show a consistent upward trend of the Gibbs free energy throughout the fetal development period, across all brain regions, culminating at birth except for the Occipital cortex areas V1C, OFC, A1C, and DFC that reach the highest Gibbs free energy at a 1-4-year point. This can be explained as the combination of dynamic cellular processes (glial mitosis versus pruning) occurring while cortexes are still developing and reach completion at around 1-2 years of age (28, 31, 32). We can associate an increase in the Gibbs free energy with a thermodynamic tendency to move out of equilibrium, which is characteristic of growth and form generation. Later in the development a dynamic equilibrium sets in whereby mitosis starts being balanced by apoptotic events that energetically stabilize the system (33) leading to a Gibbs free energy plateau.

**Figure 1.**
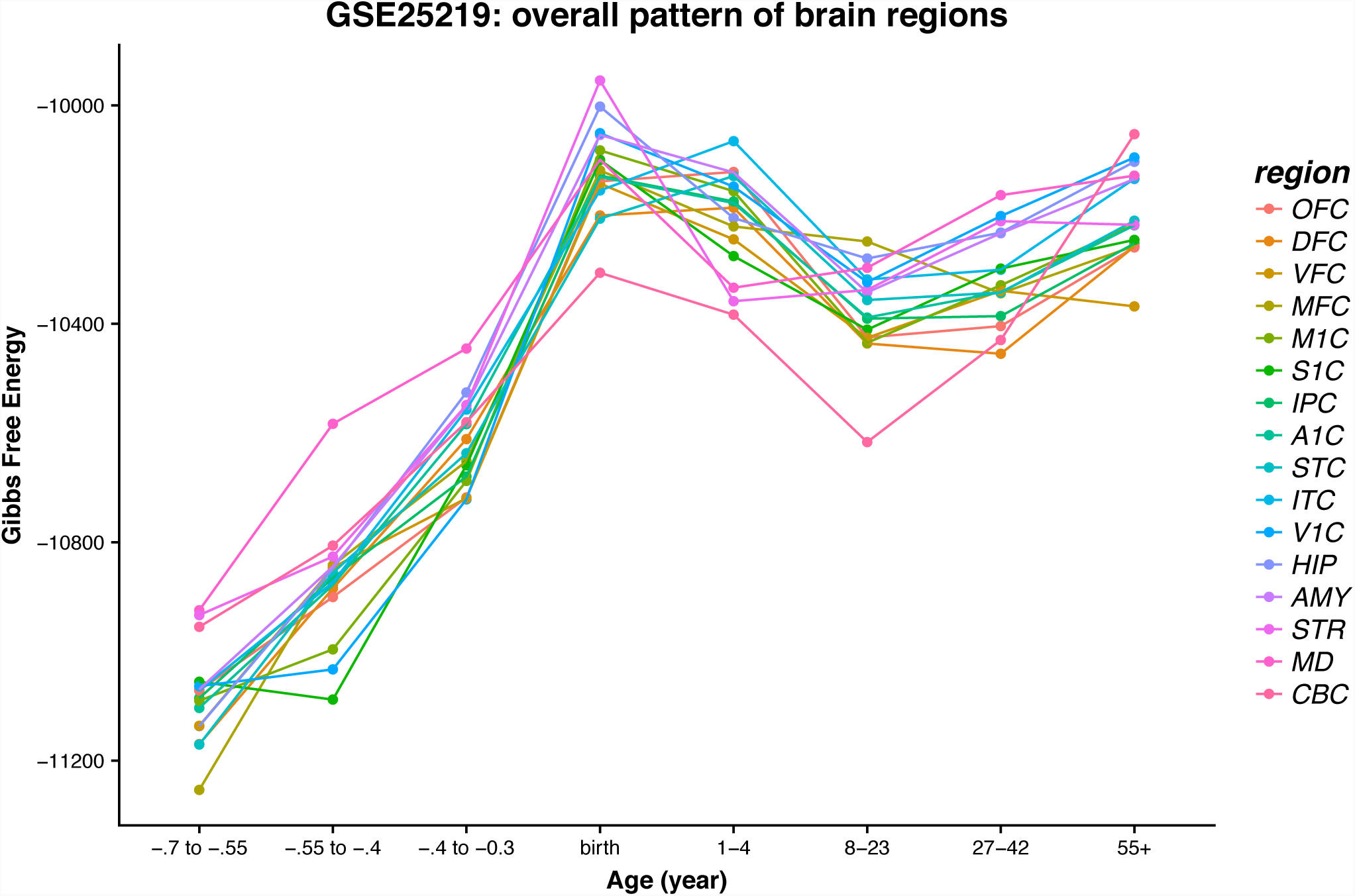
A plot of the Gibbs free energy for the 16 main brain regions averaged over the individual data sets and binned according to age groups.

At the beginning of the fetal age, the data reveal the overall lowest Gibbs free energy values, which steadily climb to a global maximum around the time of birth. This highly dynamic process taking place during the fetal development is probably due to the high rate of mitosis and physical expansion of the brain (34). Note that the Gibbs free energies are calculated based on the fetus brain specimens, ranging between 2.3 to 5.4 cm of total length (30). We hypothesize that the well-known sudden drop of the child’s body temperature at birth may be consistent with the findings of the Gibbs free energy decline and could also be responsible for reprogramming of the neuronal DNA interaction network. Change in gene interaction network levels has been demonstrated to be related to increase or decrease in entropy levels (14). Furthermore, sudden temperature change at birth appears to be the most important thermodynamic signal to neuronal DNA (35–37), hence thermodynamic does play a role at this stage.

After birth we can generally see a Gibbs free energy drop-off followed by a levelling off or slight upward trend. The data in this range are more variable between regions relative to the fetal stages. The most significant outlier we can observe is CBC, which has the lowest Gibbs free energy from childhood through adolescence and then it jumps to the highest Gibbs free energy in the 55+ range, which may be related to aging and perhaps even structural and functional deficits.

We now focus on the next stage in the evolution of the brain occurring from birth to the age of maturity, i.e. 23 years. Data in this range are more variable between brain regions relative to the fetal stages. After birth, we can generally see a Gibbs free energy drop-off followed by a leveling-off trend extended throughout childhood to sexual maturity. The brain structural/volume change, occurring after birth, is thermodynamically oriented towards its ordered developmental completion (12, 14).

In the age bracket of 8-23 years, all brain regions, except AMY and STR, show a steady decrease in their respective Gibbs free energy compared to their values at birth. This can be correlated with structural consolidation especially pronounced after reaching sexual maturity. The brain’s complete development could also send a signal to switch metabolic preferences between anabolic and catabolic energy production processes (38). Amygdala and striatum are developmentally interconnected since they both originate from the arc shaped “striatal ridge” (39–41). The singularity of the cerebellum’s structure and function has been demonstrated in the literature (42, 43). We see in our results (see Figure 1) that CBC reaches the lowest Gibbs free energy value in the age bracket from 8 to 23 years when cerebellum is known to reach its complete development. Immediately after, MRI studies show a marginal volume decline that at the age of 50 becomes exponential, which could explain our findings (44–46).

### Comparisons Between Male and Female Data Sets

In this section we specifically analyze the differences between the Gibbs free energy results obtained for the male and female brains, respectively. We discuss these differences for each area of the brain separately.

For the **primary auditory cortex** A1C (see Figure 2), for each age group there is no difference between male and female subjects beyond statistical error. A significant increase in Gibbs free energy can be observed, peaking at birth and then dropping moderately until the adolescent age region (ages 8-23) is reached, and then increasing slightly again for the last two age brackets. Generally, changes appear more exaggerated for the female subjects (lower troughs and higher peaks in the Gibbs free energy values).

**Figure 2.**
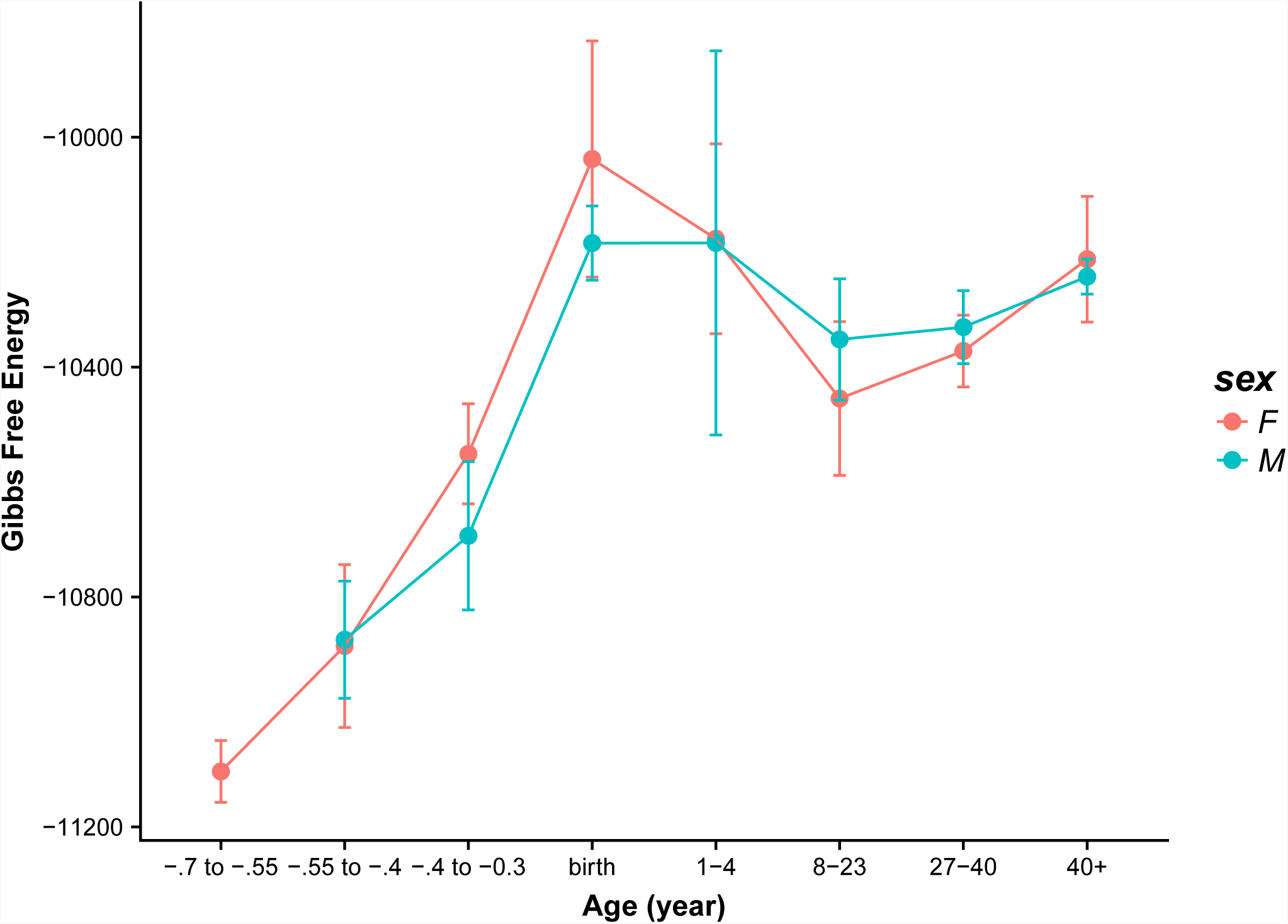
Gibbs free energy for male and female samples in the primary auditory (A1C) cortex.

For the **amygdala** AMY (see Figure 3), the overall pattern is as described for A1C, above. The female regions are consistently greater than the male counterparts until birth at which point the Gibbs free energy values for female samples peak. The corresponding values for male samples peak in the next bracket (1-4), however there is a significant error associated with this data point. This is followed by a dip in the adolescent bracket and a slight increase after. Again, we see a crossover along the age axis between the male and female Gibbs free energy values; the male subjects are characterized by more stable Gibbs free energy profiles, particularly in the last three age brackets (8+).

**Figure 3.**
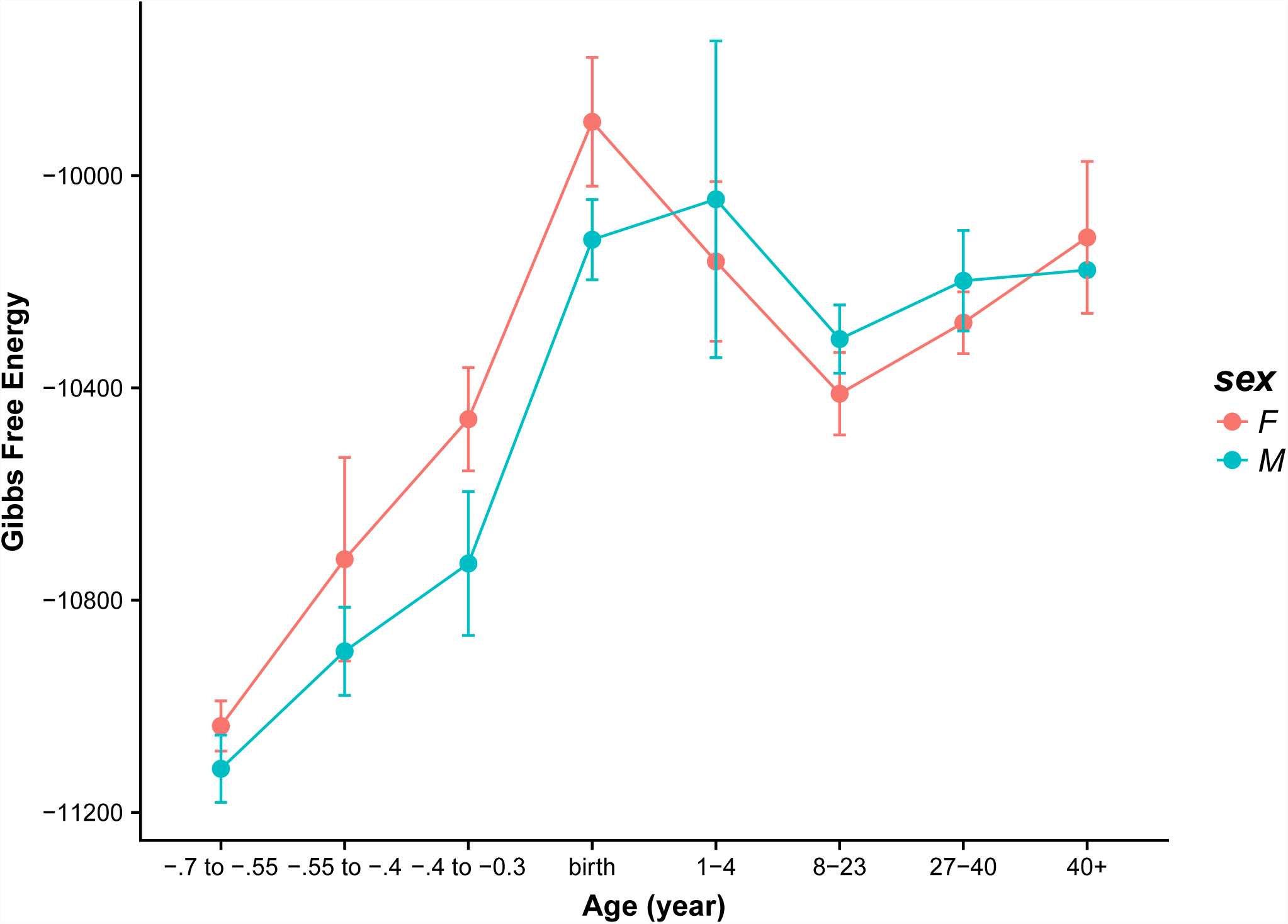
Gibbs free energy for male and female samples in the amygdala (AMY).

The data for the **cerebellar cortex** CBC (see Figure 4) seem to differ relative to many of the sex plots. Here, the fetal region does not have as significant of an upward trend. For the female subjects there is a moderate upward trend and then a sudden peak at birth that immediately drops again after birth. The male region does not appear to have the typical linear upward slope, however there is no data for the youngest range and a relatively large statistical error in the second range. At birth, the male samples show Gibbs free energies, which are significantly lower than those for the females. After birth, the male and female samples effectively move in unison with no statistical differences. The dip and upward swing after age 8 is more pronounced for the cerebellar cortex

**Figure 4.**
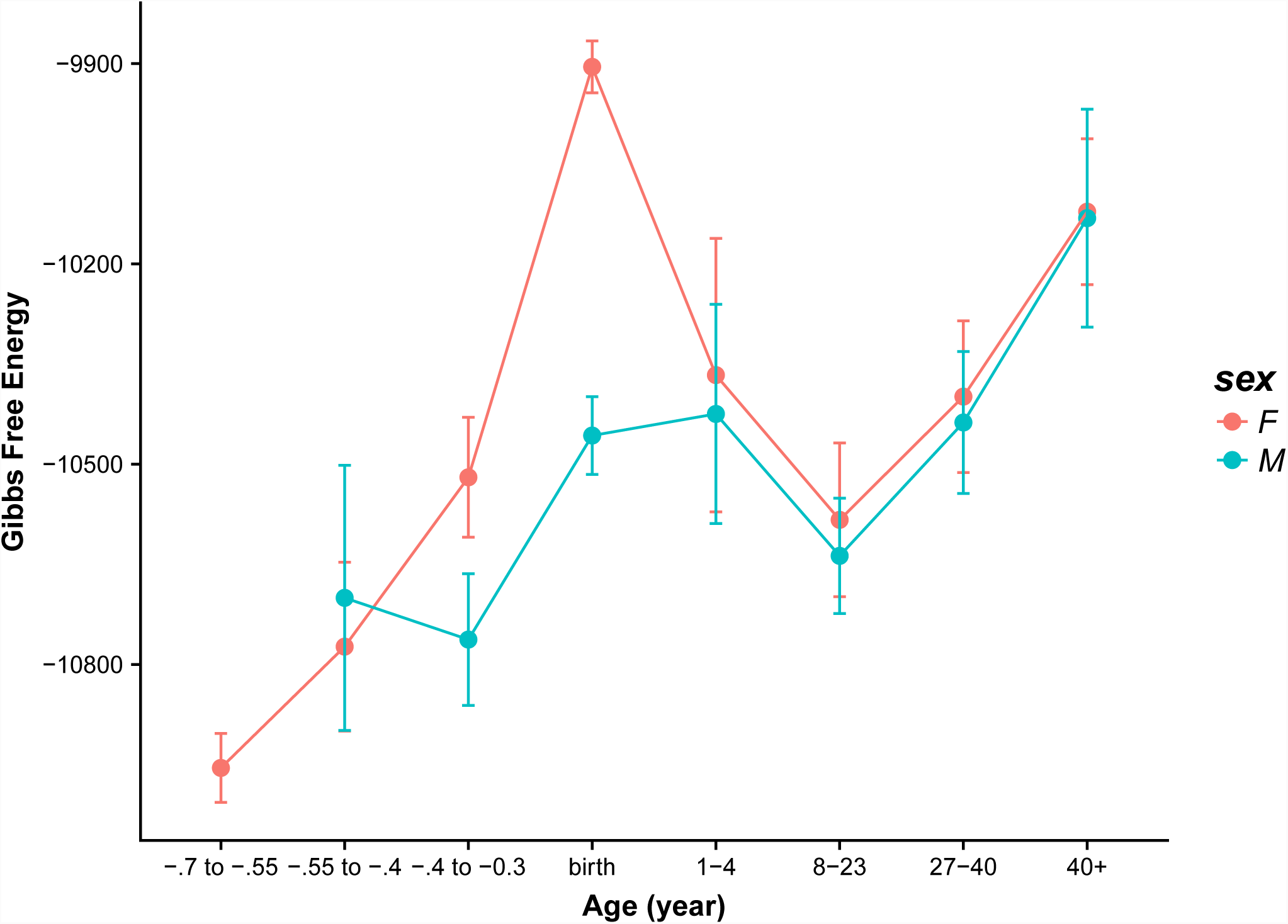
Gibbs free energy for male and female samples in the cerebellar cortex (CBC).

The data for the **dorsolateral prefrontal cortex** DFC seems to fall somewhere between the AMY and A1C, and CBC data (see Figure 5). Generally, there is an upward trend in the Gibbs free energy from the inception of the fetal stage until birth. Similarly to the CBC data, the upward trend is not present between the second and third age bracket (covers −0.55 to −0.3) for the male subjects only. Again, the male subject Gibbs free energy peaks slightly over birth in the 1-4 bracket. Both sexes exhibit the 8+ dip and swing. As with A1C and AMY, this dip and swing is more pronounced for female subjects.

**Figure 5.**
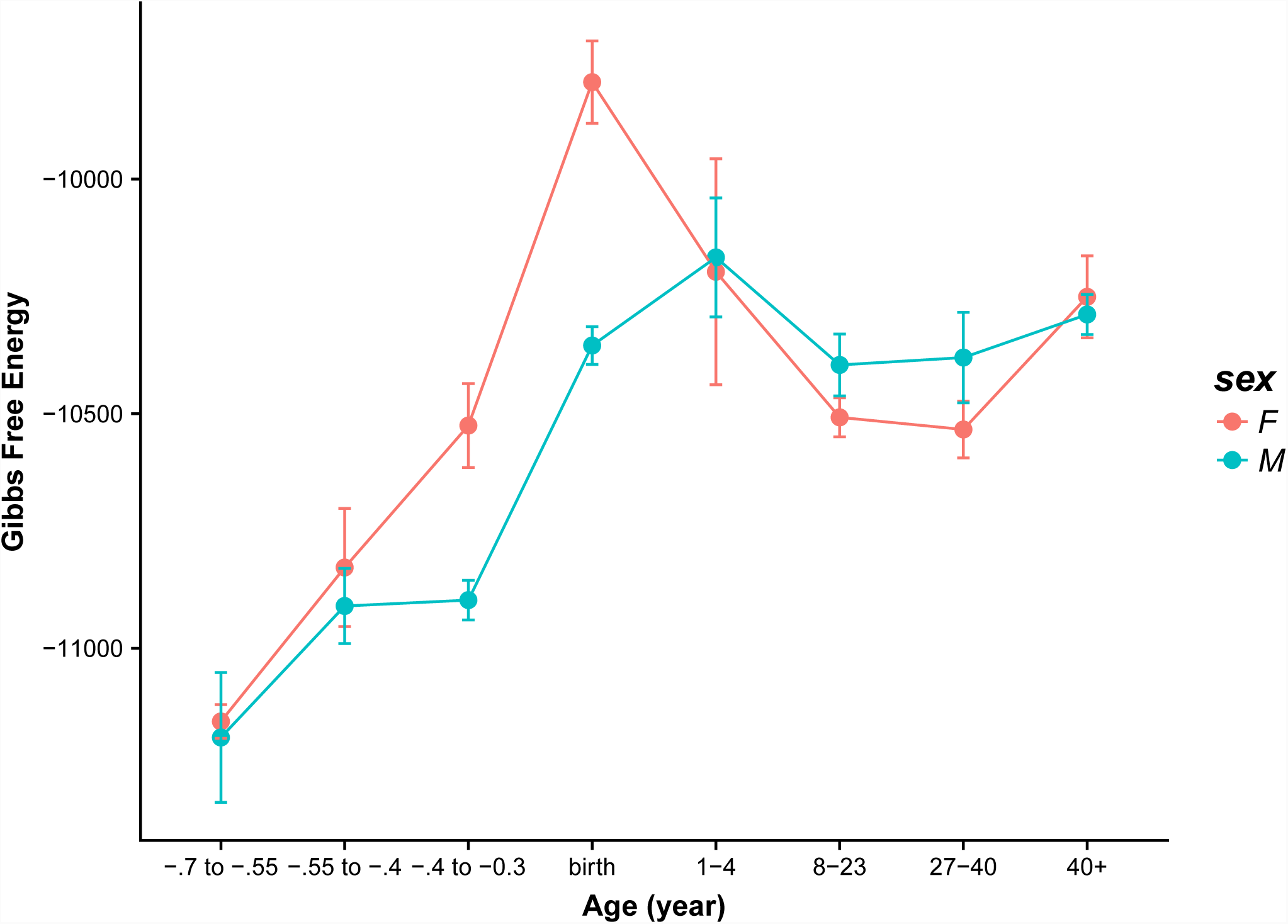
Gibbs free energy for male and female samples in the dorsolateral prefrontal cortex (DFC).

In the **hippocampus** HIP data (see Figure 6), we observe the same upward trends until birth. Overall, the data for the male and female regions appear quite similar. Both have a peak at birth (though more significant for the female region). Unlike other regions, there, the Gibbs free energy is more stable after birth, particularly for the male regions. The female subjects again have an upward swing in the Gibbs free energy but mostly in the 40+years region.

**Figure 6.**
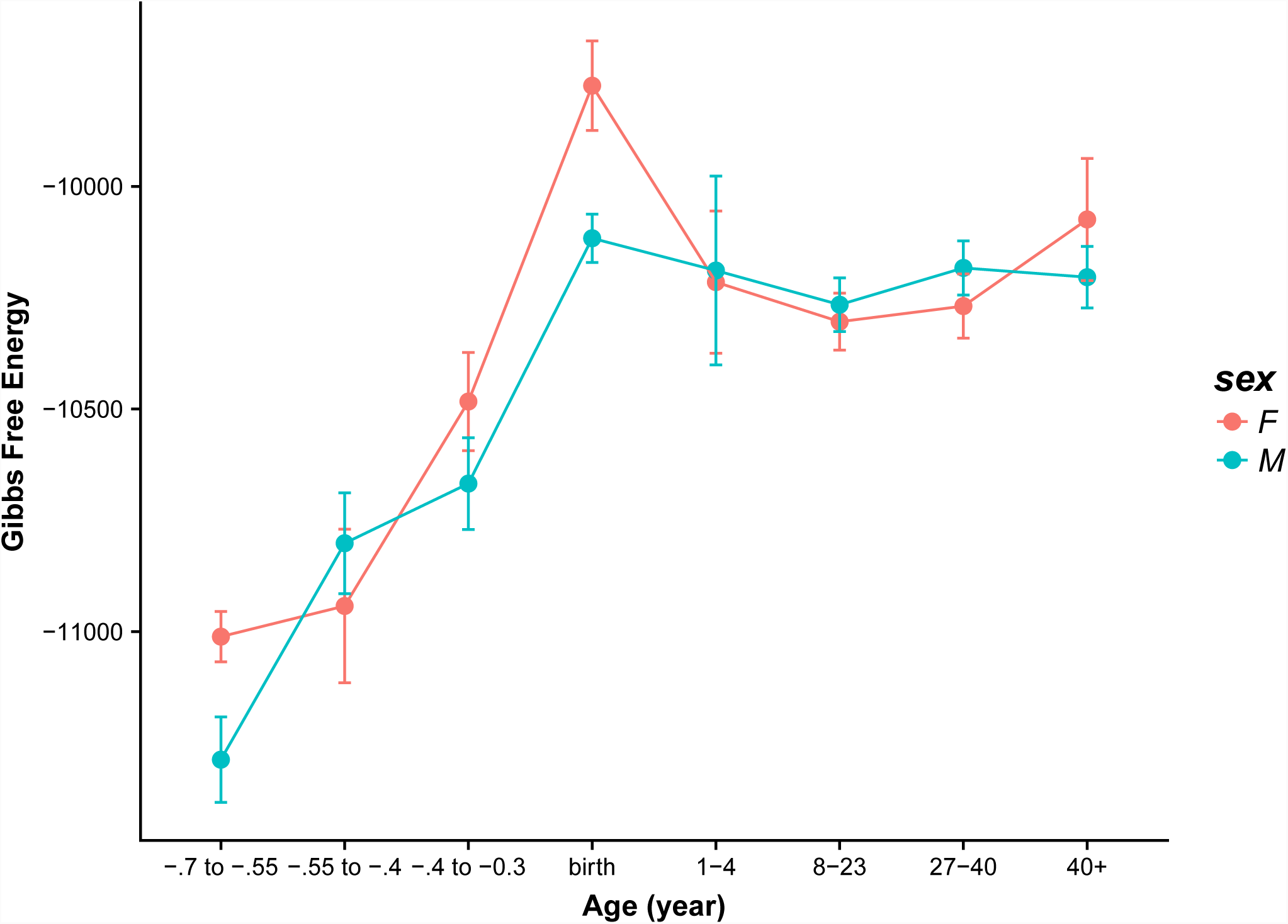
Gibbs free energy for male and female samples in the hippocampus (HIP).

For the data for the **posterior inferior parietal cortex** IPC (see Figure 7), the upward trend is more modest in the fetal samples. The female samples have a sudden jump at birth and then the Gibbs free energy drops down abruptly (although not to fetal levels). The male subjects also show a large increase at birth but this continues into the 1-4 age bracket (however, there is significant error associated with this data point). The typical dip and swing is then observed as well as the sex crossover mentioned above.

**Figure 7.**
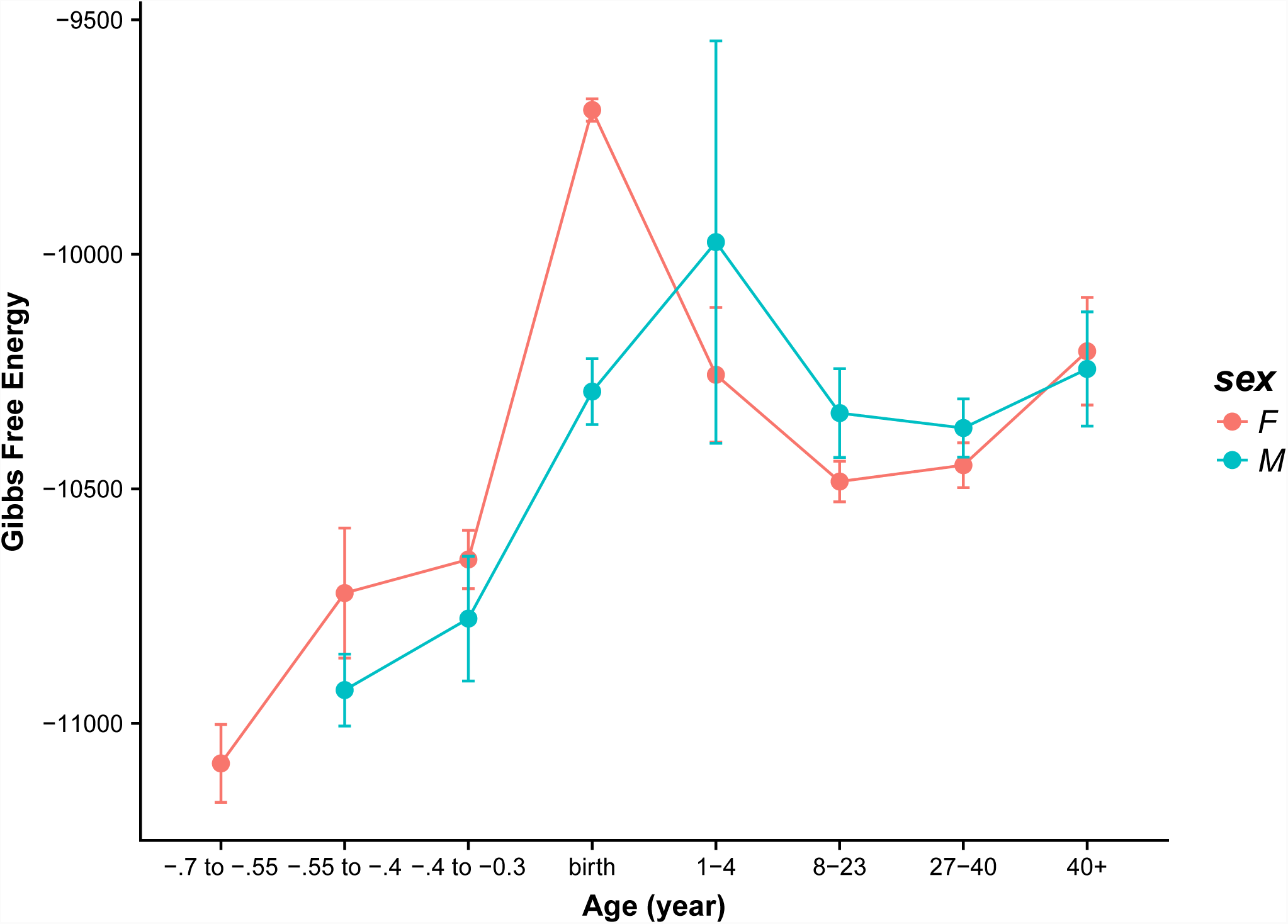
Gibbs free energy for male and female samples in the posterior inferior parietal cortex (IPC).

For the fetal period for the **inferior temporal cortex** ITC (see Figure 8), we can observe an upward slope in the Gibbs free energy, however its magnitude is greater for the female subjects. The female regions exhibit the typical peak at birth, followed by a dip and then increase again at age 40+. The male region has a clear peak in the 1-4 age range, then it dips modestly, and then remains relatively stable. Again, we can see a sex crossover effect after birth.

**Figure 8.**
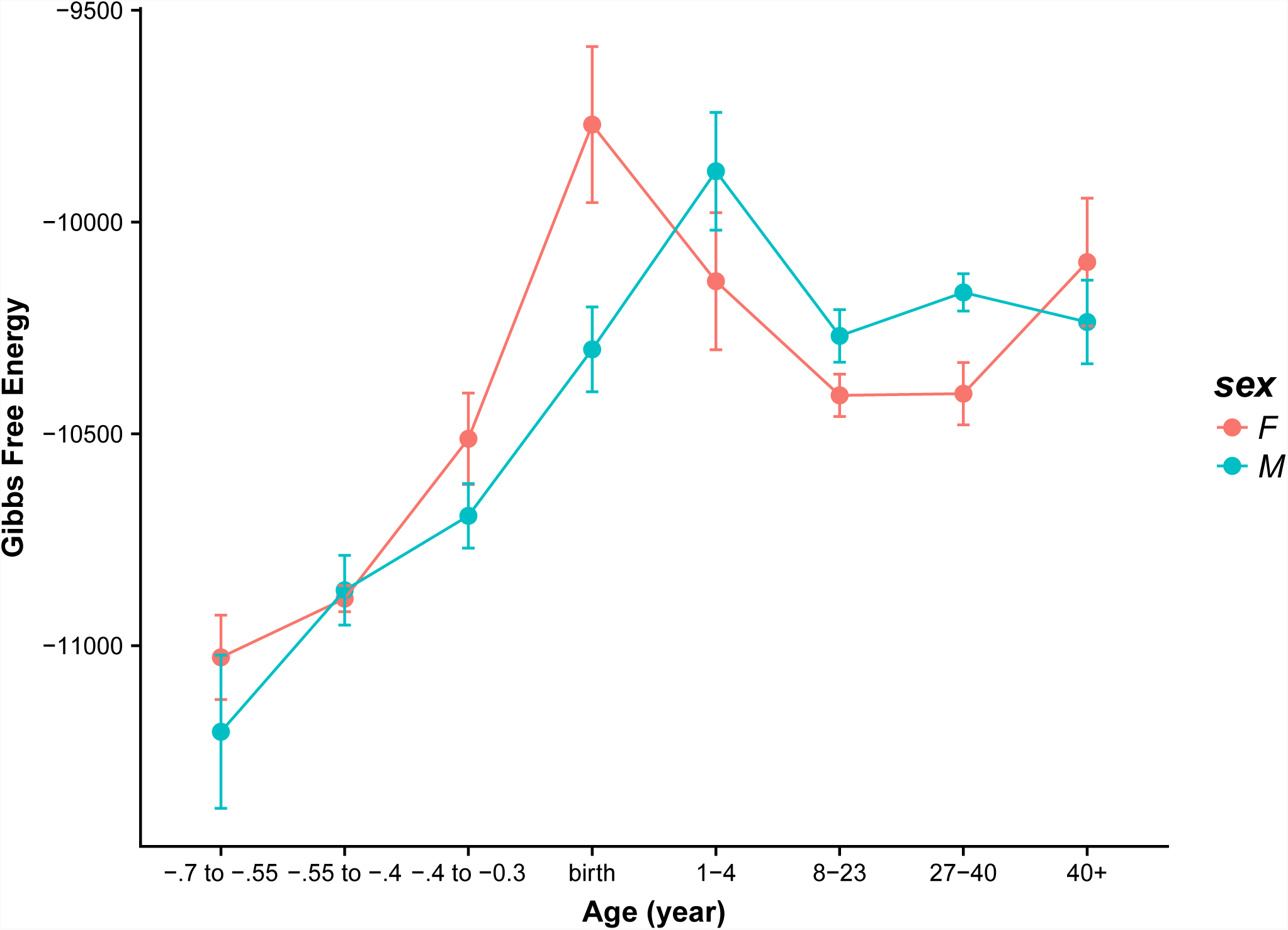
Gibbs free energy for male and female samples in the inferior temporal cortex (ITC).

The Gibbs values for the **primary motor cortex** M1C (see Figure 9) are quite typical relative to other brain regions. There is an upward trend, peaking at birth for the female samples and peaking slightly in the 1-4 region for the male subjects. Both sexes move approximately in unison, dipping in the 8-23 bracket and then swing upward slightly thereafter.

**Figure 9.**
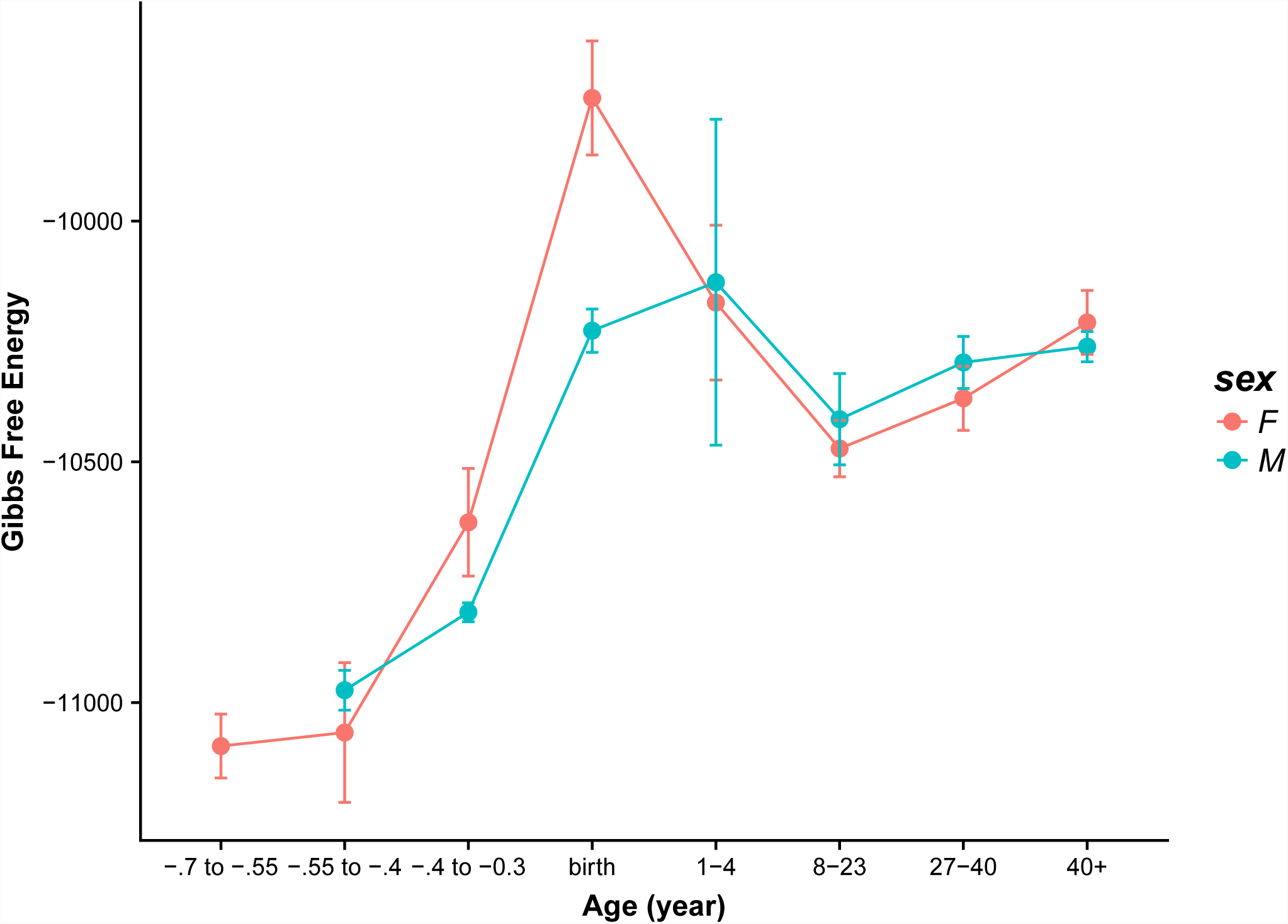
Gibbs free energy for male and female samples in the primary motor (M1C) cortex.

The Gibbs values for the **mediodorsal nucleus** of the thalamus MD (see Figure 10) are somewhat atypical relative to other brain regions. On reason for this may be that there was less data for this region (in particular it is worth noting that there was only one male sample in the 1-4 age bracket). The female subjects follow an upward trend, peak at birth and then dip down after birth and remain relatively stable. The male curve is more sporadic, peaking at birth then dropping off abruptly and swing back up to approximately the same value observed at birth.

**Figure 10.**
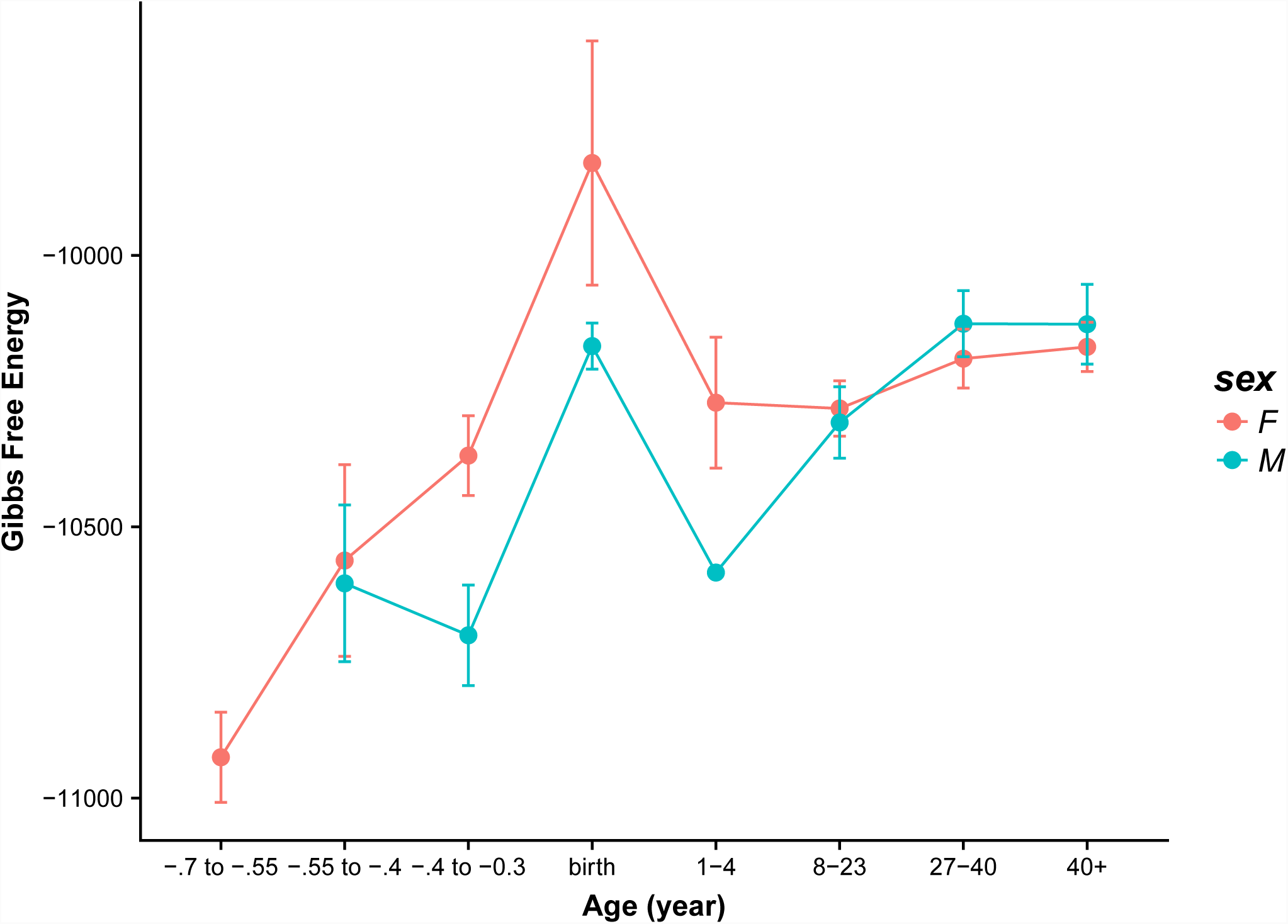
Gibbs free energy for male and female samples in the mediodorsal nucleus (MD) of thalamus.

For the **medial prefrontal cortex** MFC (see Figure 11), we see a typical pattern in the female region: an increase during fetal development, a peak at birth immediately followed by a dip, and then slight upward swing in the 40+ region. The male regions on the hand, follow a similar fetal developmental pattern (translated down slightly), however the is no dip and swing. Instead we basically see a plateau after birth and a slight drop in the 40+ region.

**Figure 11.**
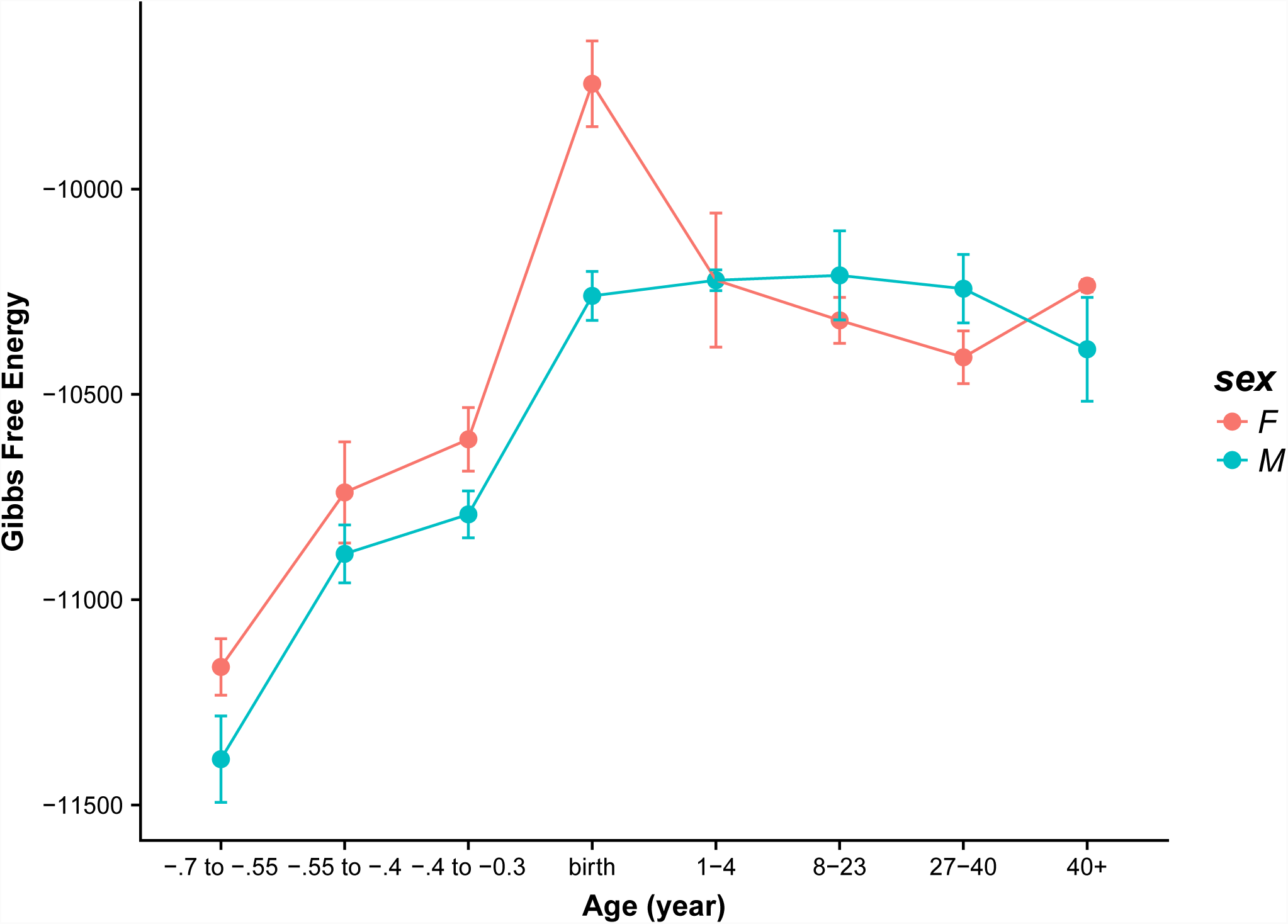
Gibbs free energy for male and female samples in the medial prefrontal cortex (MFC).

The **orbital prefrontal cortex** OFC (see Figure 12) follows a pattern similar to the one observed in the ITC (see Figure 8). The difference here is that the jump between the third fetal period and birth is more significant than in the ITC. Like the ITC the male region peaks after birth (however there is only one data point) and is stable after it dips down in the 8-23 range. The female curve follows the dip and swing and we also see the sex crossover pattern.

**Figure 12.**
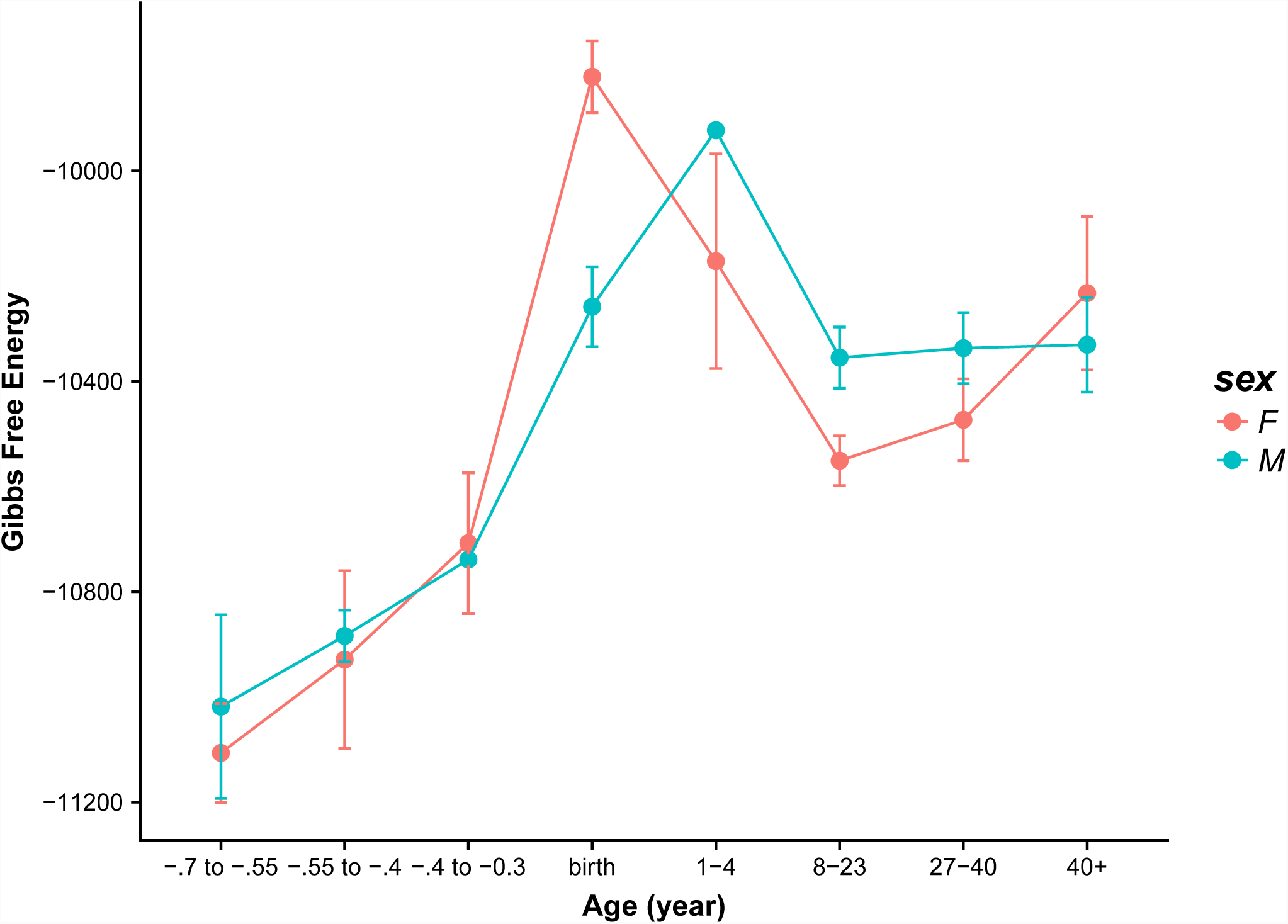
Gibbs free energy for male and female samples in the orbital prefrontal cortex (OFC).

The Gibbs values for the **primary somatosensory cortex** S1C (see Figure 13) again display an upward trend during fetal development and a peak at birth for the female subjects, with the male values increasing slightly after birth. Both curves dip into the 8-23 age bracket however the male region has an anomalous peak again in the 27-40 age bracket. The female region remains relatively stable after the dip.

**Figure 13.**
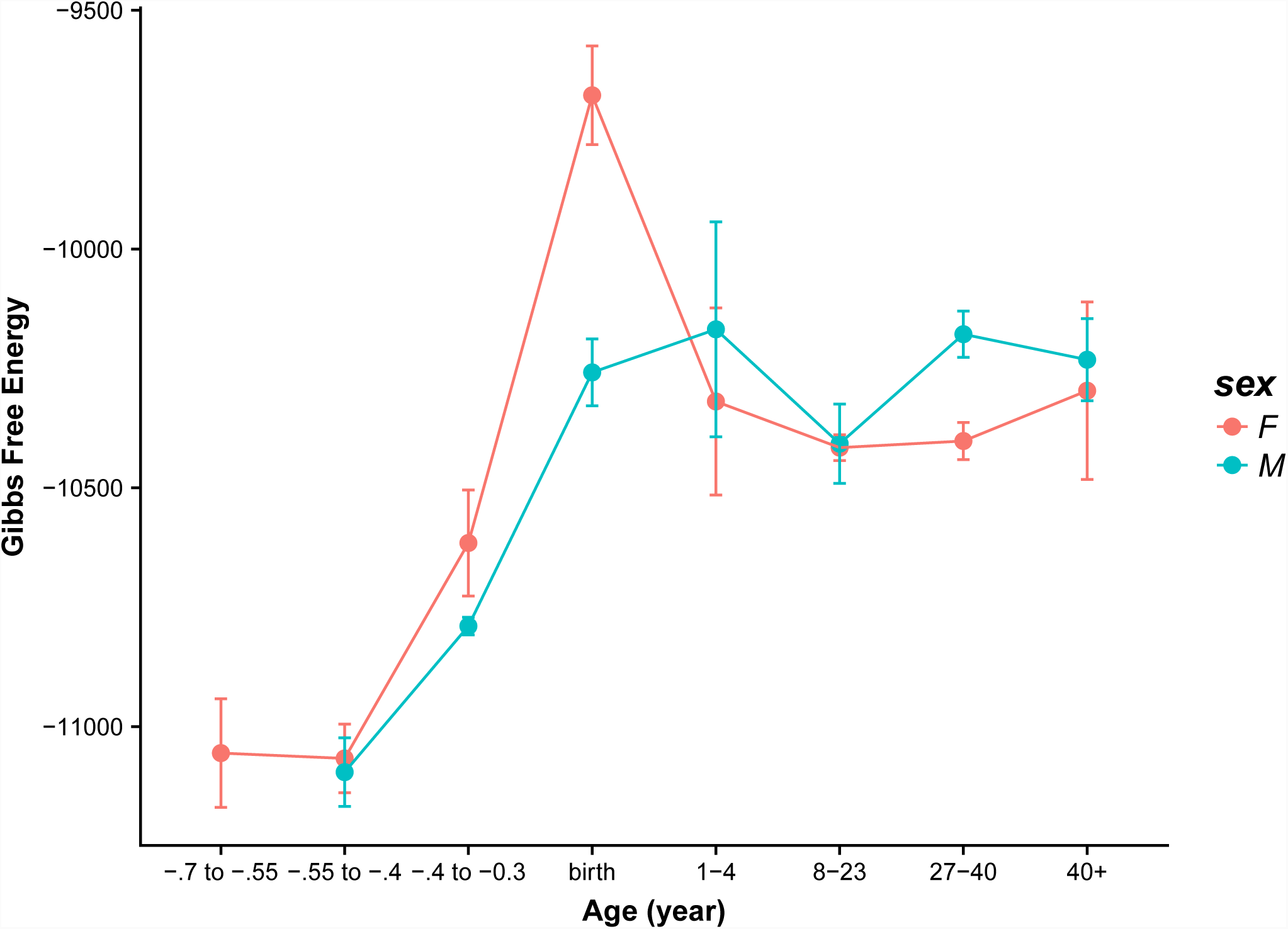
Gibbs free energy for male and female samples in the primary somatosensory (S1C) cortex.

The **posterior superior temporal cortex** STC (see Figure 14) contains many of the patterns observed in the other brain regions. One thing that makes this region unique, is that it is the only one where the highest male value is greater than the largest female Gibbs energy. This may in part be due to the female peak being a bit less pronounced. Overall, we see the upward trend during fetal development (with a stall again between the second and third period for male samples), with peaks at birth and in the 1-4 age bracket for female and male samples, respectively. The female region has the dip and swing whereas the male samples are relatively flat in later years. The pattern also seems reminiscent of V1C (see Figure 16).

**Figure 14.**
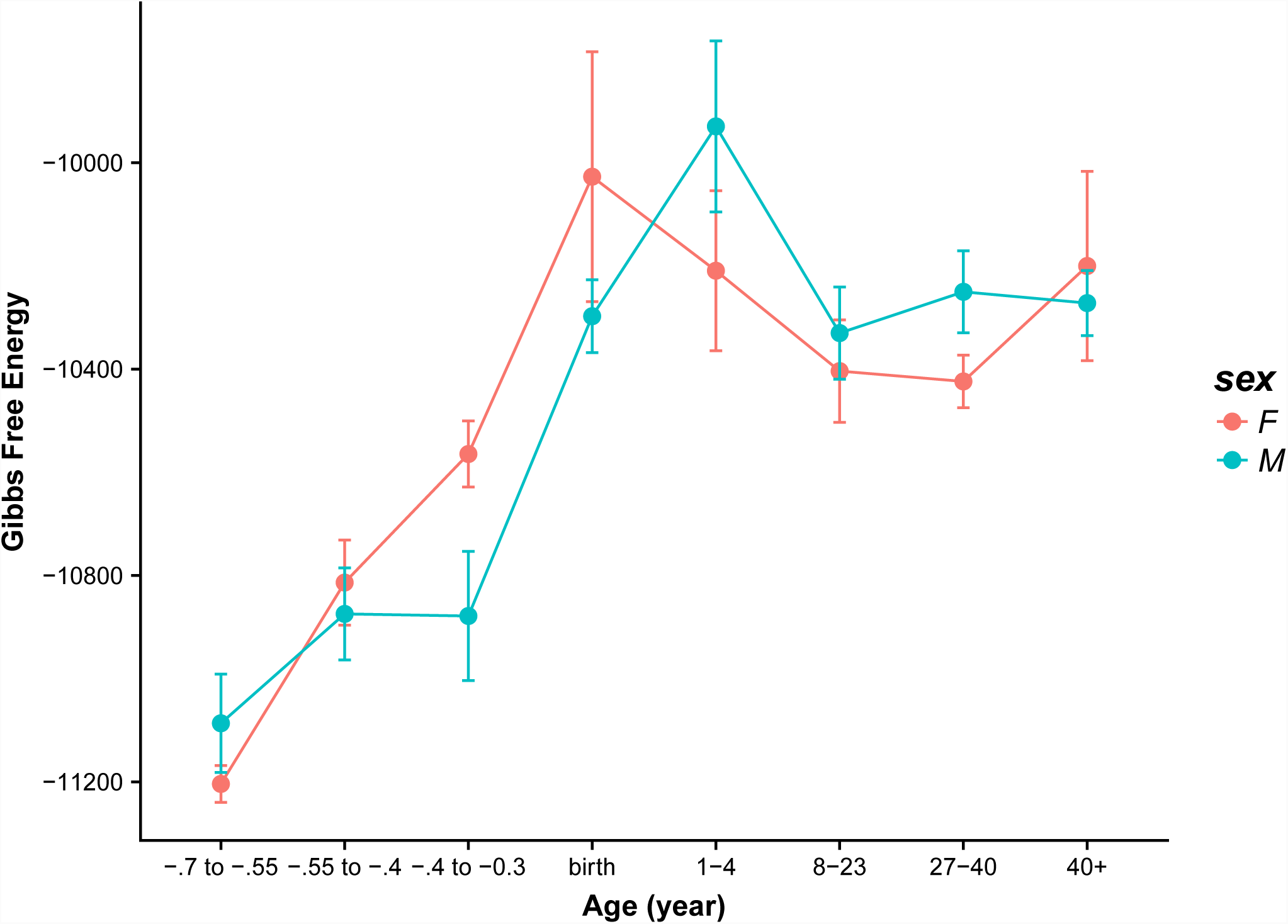
Gibbs free energy for male and female samples in the posterior superior temporal cortex (STC).

For the **striatum** STR (see Figure 15), we see somewhat erratic behaviour similar to that which was observed in the MD (see Figure 10). Like the MD, this may in part be due to fewer samples being available in this region (note, only one male sample in the −0.4 to −0.3 and 1-4 age brackets). The female region, like in the MD follows an upward trend until peaking at birth. The curve then dips down during the toddler years but then remains stable thereafter.

**Figure 15.**
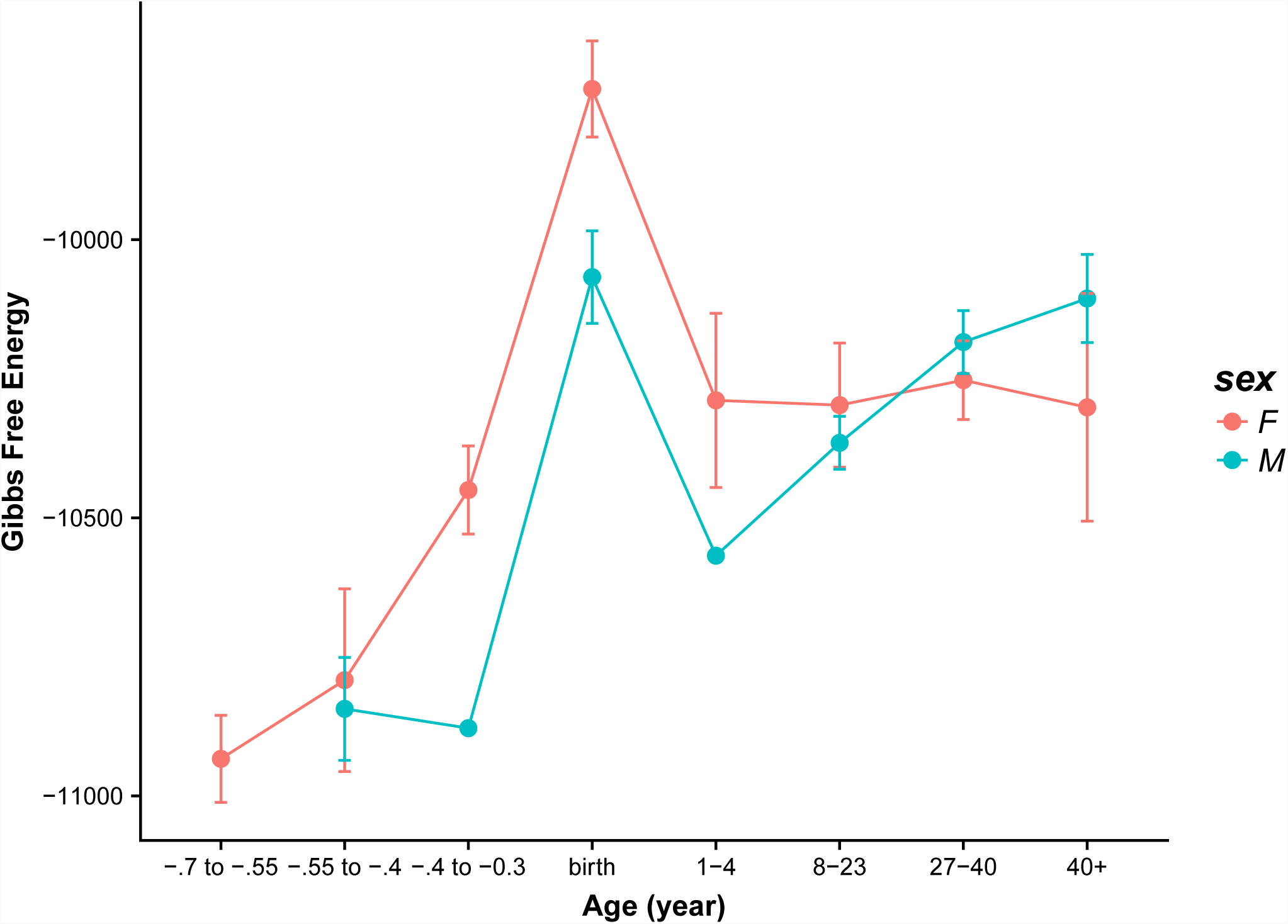
Gibbs free energy for male and female samples in the striatum (STR).

The female samples for the **primary visual cortex** V1C (see Figure 16) follow a typical pattern: upward slope during fetal development, peak at birth, dipping until 8-23 bracket, followed by an upward swing. The male pattern is similar to that observed in the S1C (see Figure 12): peaking after birth, then dipping in the 8-23 bracket and increasing again, plateauing from age 27 onwards. Overall, V1C and S1C appear quite similar.

**Figure 16.**
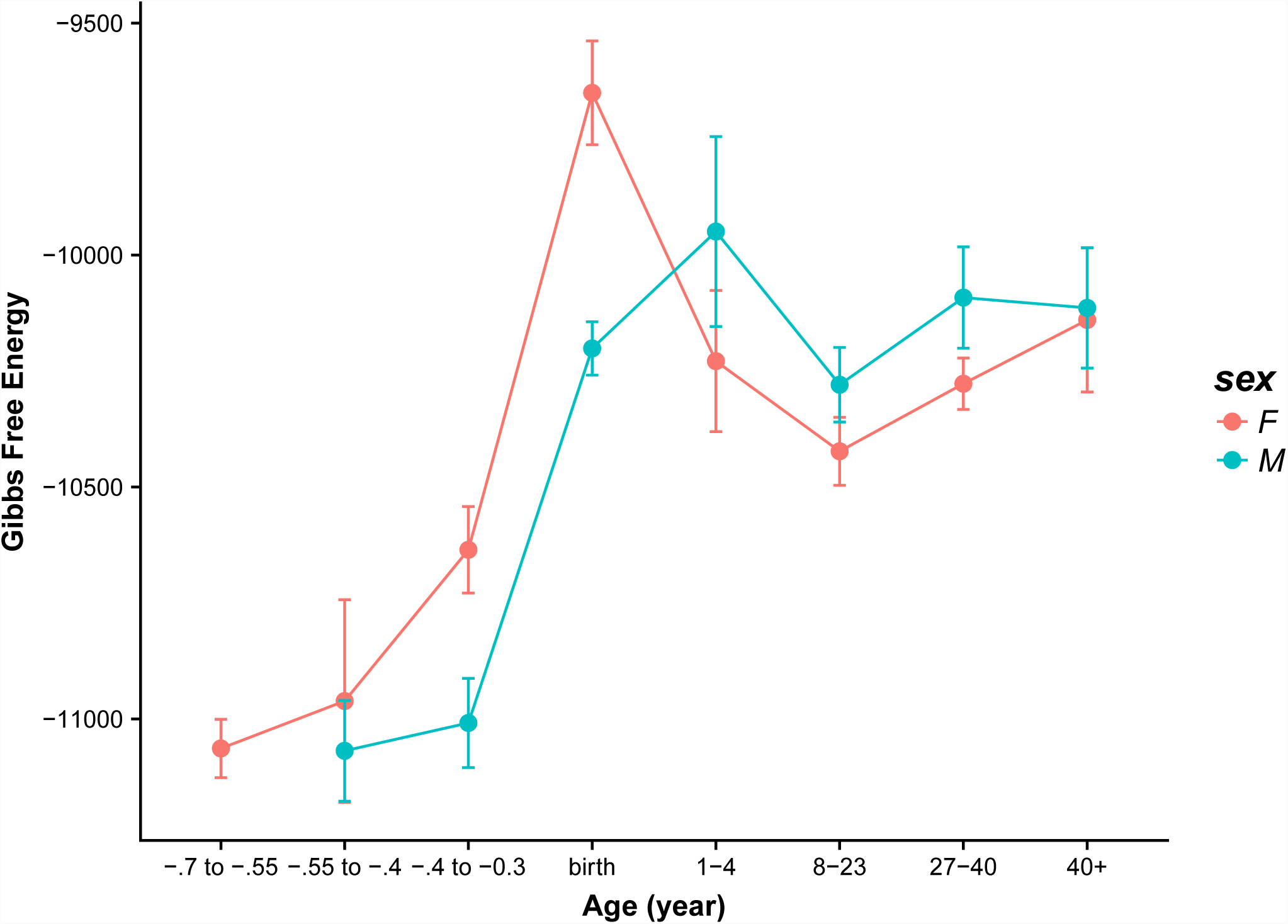
Gibbs free energy for male and female samples in the primary visual (V1C) cortex.

Overall, there seems to be more stability in the **ventrolateral prefrontal cortex** VFC (see Figure 17) relative to other brain regions. The female samples show only a slight increase in the Gibbs free energy during fetal development and then a jump at birth. The Gibbs free energy drops linearly until the 8-23 bracket after which point the curve is effectively flat. For the male subjects, there is a step from the first fetal period to the second and then another step at birth, after which point the Gibbs free energy values remain relatively unchanged. To demonstrate that the cross-over effect is a general effect and not limited to specific regions of the brain, we have averaged the Gibbs free energy values over all brain regions for males and females separately and plotted the results in Fig. 18. As can be readily seen, the cross-over effect in early childhood, around the age 1-4 years is consistently present. Moreover, in middle age, around the age of 40, both female and male brains converge to the same value of the Gibbs free energy.

**Figure 17.**
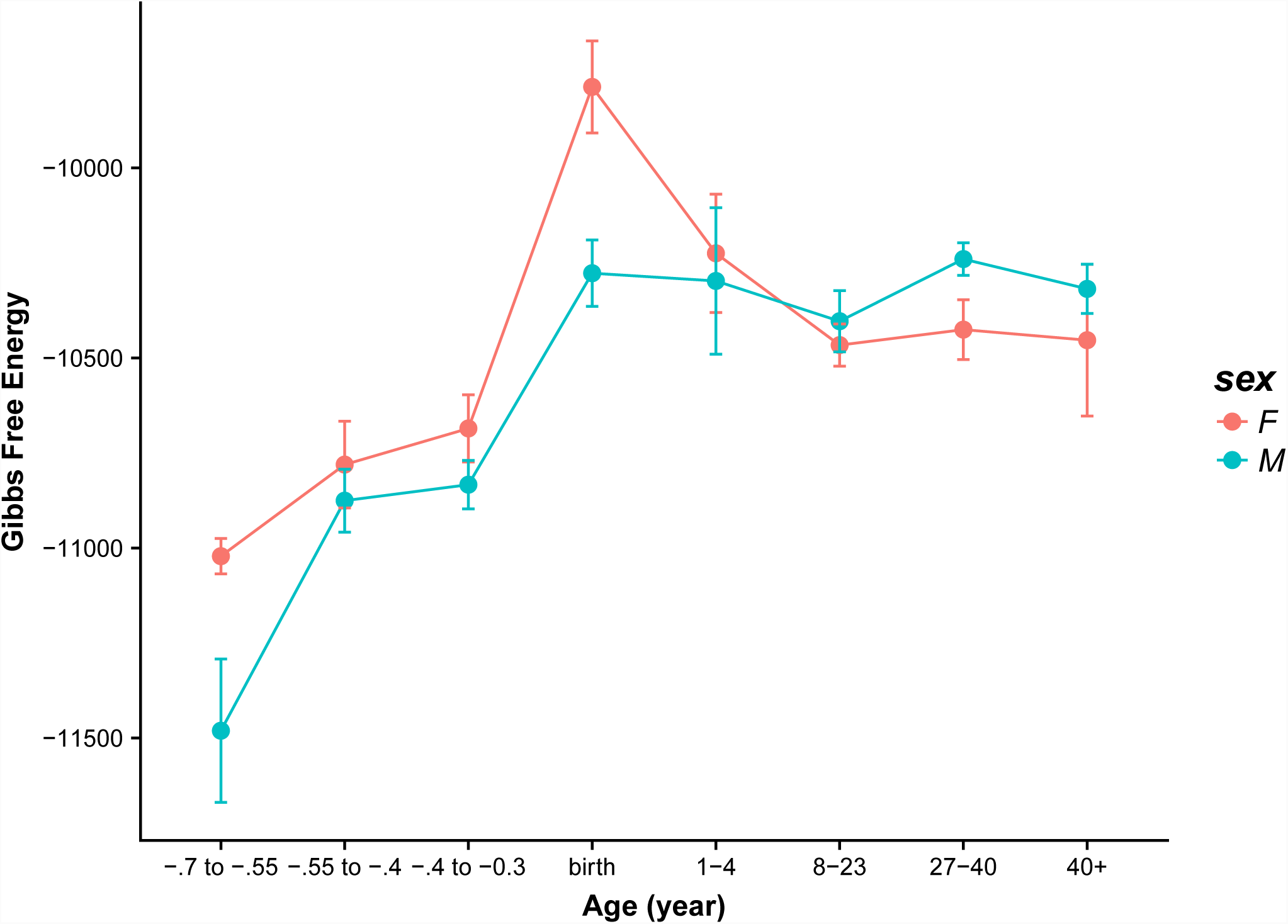
Gibbs free energy for male and female samples in the ventrolateral prefrontal cortex (VFC).

**Figure 18.**
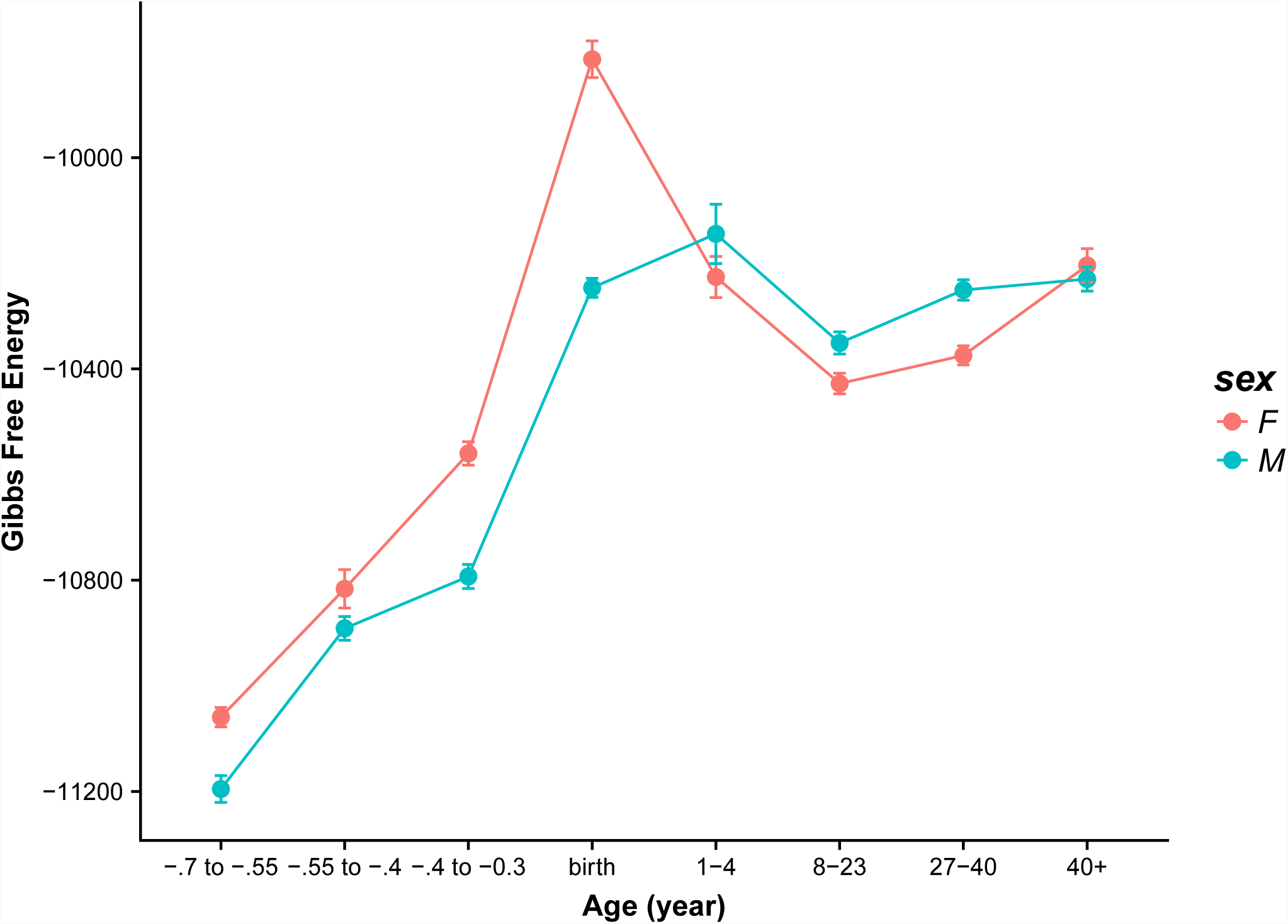
Plot of the Gibbs free energy values averaged over all 16 brain areas and presented for female and male cases separately.

## Discussion

In this paper we have analyzed protein expression data obtained from biopsied brains of females and males spanning a range of ages from prenatal to old age. The data were organized according to their origin involving 16 distinct and well-defined brain regions. We have introduced a quantitative thermodynamic measure that reflects the structural brain development level based on the protein-protein interaction network’s characteristics. This measure is the network’s Gibbs free energy, which originates in statistical physics where its increase signifies departure from a thermodynamic equilibrium. Conversely, a tendency of the thermodynamic system to attain the lowest Gibbs free energy under existing constraints is a consequence of the second law of thermodynamics and is consistent a maximum entropy principle. Here, we applied this measure to address the question if there are any trends that correlate with age and secondly if there are differences in this regard between males and females. Our analysis shows a dramatic increase in the Gibbs free energy in prenatal brains as they approach the time of birth at which point the Gibbs free energy values consistently achieve a maximum value possibly reflecting the brain’s structural and functional development. Subsequently, the Gibbs free energy decreases with age achieving a local minimum around the age of sexual maturity. From that age onward, the Gibbs free energy slowly increases in almost all brain regions. While we offer no concrete explanation of this effect, we hypothesize that this may reflect the brain’s synaptic plasticity due to continued learning and accumulation of life’s experience. Finally, we have also compared the Gibbs free energy for male and female samples, respectively. In most brain regions and also for the averaged values over all regions of the brain we found that from prenatal stages of development until early childhood the Gibbs free energy from female subjects is higher than that of the male counterparts, which possibly signifies a more rapid development of female brains than is the case for male brains. A cross-over effect can be seen in early childhood where the relationship is reversed but eventually, around the age of 40, both curves converge to the same value. Admittedly, the relatively small sample size suggests that these results should be interpreted with great caution. Larger sets of similar data would offer greater confidence in our analysis but due to lack of such datasets we are unable to provide more statistically significant conclusions.

## Author Contributions

EAR and JAT conceived of the project. ST did the analysis. HTS assisted in the analysis. MC helped interpret the results. All authors contributed to writing the manuscript.

## Acknowledgments

We gratefully acknowledge Philip Winter for programming and editorial assistance. EAR acknowledges funding from Office of Naval Research (ONR)/N00014-15-2126. The content is solely the responsibility of the authors and does not necessarily represent the official views of the US Navy. EAR also acknowledges partial funding from CSTS-Healthcare, Toronto, Canada. JAT gratefully acknowledges research support from the Natural Sciences and Engineering Research Council of Canada.

## References

1. McGuire SH, Rietman EA, Siegelmann H, Tuszynski JA (2017) Gibbs free energy as a measure of complexity correlates with time within C. elegans embryonic development. J Biol Phys 43(4):551–563.

2. Breitkreutz D, Hlatky L, Rietman E, Tuszynski JA (2012) Molecular signaling network complexity is correlated with cancer patient survivability. Proc Natl Acad Sci U S A 109(23):9209–9212.

3. Hinow P, Rietman EA, Omar SI, Tuszyński JA (2015) Algebraic and topological indices of molecular pathway networks in human cancers. Math Biosci Eng MBE 12(6):1289–1302.

4. Benzekry S, Tuszynski JA, Rietman EA, Lakka Klement G (2015) Design principles for cancer therapy guided by changes in complexity of protein-protein interaction networks. Biol Direct 10:32.

5. Rietman EA, Platig J, Tuszynski JA, Lakka Klement G (2016) Thermodynamic measures of cancer: Gibbs free energy and entropy of protein-protein interactions. J Biol Phys 42(3):339–350.

6. Rietman EA, Scott JG, Tuszynski JA, Klement GL (2017) Personalized anticancer therapy selection using molecular landscape topology and thermodynamics. Oncotarget 8(12):18735–18745.

7. Huang S (2011) On the intrinsic inevitability of cancer: from foetal to fatal attraction. Semin Cancer Biol 21(3):183–199.

8. Moris N, Pina C, Arias AM (2016) Transition states and cell fate decisions in epigenetic landscapes. Nat Rev Genet 17(11):693–703.

9. Uhlén M, et al. (2015) Proteomics. Tissue-based map of the human proteome. Science 347(6220):1260419.

10. The Human Protein Atlas (2018) Available at: https://www.proteinatlas.org/ [Accessed December 19, 2018].

11. Sjöstedt E, et al. (2015) Defining the Human Brain Proteome Using Transcriptomics and Antibody-Based Profiling with a Focus on the Cerebral Cortex. PloS One 10(6):e0130028.

12. Kang HJ, et al. (2011) Spatio-temporal transcriptome of the human brain. Nature 478(7370):483–489.

13. Aitchison JD, Rout MP (2015) The interactome challenge. J Cell Biol 211(4):729–732.

14. Yao Y, et al. (2013) The increase of the functional entropy of the human brain with age. Sci Rep 3:2853.

15. Kim M-S, et al. (2014) A draft map of the human proteome. Nature 509(7502):575–581.

16. Wilhelm M, et al. (2014) Mass-spectrometry-based draft of the human proteome. Nature 509(7502):582–587.

17. Guo Y, et al. (2013) Large scale comparison of gene expression levels by microarrays and RNAseq using TCGA data. PloS One 8(8):e71462.

18. TCGA: The Cancer Genome Atlas (2018) Available at: https://cancergenome.nih.gov/ [Accessed December 20, 2018].

19. Chatr-Aryamontri A, et al. (2017) The BioGRID interaction database: 2017 update. Nucleic Acids Res 45(D1):D369–D379.

20. BioGRID (2018) Available at: https://thebiogrid.org/ [Accessed December 20, 2018].

21. Sedmak G, et al. (2016) Developmental Expression Patterns of KCC2 and Functionally Associated Molecules in the Human Brain. Cereb Cortex N Y N 1991 26(12):4574–4589.

22. Edgar R, Domrachev M, Lash AE (2002) Gene Expression Omnibus: NCBI gene expression and hybridization array data repository. Nucleic Acids Res 30(1):207–210.

23. GEO: Gene Expression Omnibus (2018) Available at: https://www.ncbi.nlm.nih.gov/geo/ [Accessed December 20, 2018].

24. Colantuoni C, et al. (2011) Temporal dynamics and genetic control of transcription in the human prefrontal cortex. Nature 478(7370):519–523.

25. Somel M, et al. (2010) MicroRNA, mRNA, and protein expression link development and aging in human and macaque brain. Genome Res 20(9):1207–1218.

26. Miller JA, et al. (2014) Transcriptional landscape of the prenatal human brain. Nature 508(7495):199–206.

27. BrainSpan: Atlas of the Developing Human Brain (2018) Available at: http://www.brainspan.org/ [Accessed December 20, 2018].

28. Stiles J, Jernigan TL (2010) The basics of brain development. Neuropsychol Rev 20(4):327–348.

29. Blackshaw S, et al. (2010) Molecular pathways controlling development of thalamus and hypothalamus: from neural specification to circuit formation. J Neurosci Off J Soc Neurosci 30(45):14925–14930.

30. Hill MA (2018) Embryology - Timeline human development. Available at: https://embryology.med.unsw.edu.au/embryology/index.php/Timeline_human_development [Accessed December 19, 2018].

31. Budday S, Steinmann P, Kuhl E (2015) Physical biology of human brain development. Front Cell Neurosci 9:257.

32. Moura da Costa e Sousa AM (2013) Human brain development and evolution: insights from gene expression. Available at: https://repositorio-aberto.up.pt/handle/10216/76388 [Accessed December 19, 2018].

33. Kim Y, Han S, Choi S, Hwang D (2014) Inference of dynamic networks using time-course data. Brief Bioinform 15(2):212–228.

34. Luecke RH, Wosilait WD, Young JF (1999) Mathematical modeling of human embryonic and fetal growth rates. Growth Dev Aging GDA 63(1–2):49–59.

35. Morton SU, Brodsky D (2016) Fetal Physiology and the Transition to Extrauterine Life. Clin Perinatol 43(3):395–407.

36. Asakura H (2004) Fetal and neonatal thermoregulation. J Nippon Med Sch Nippon Ika Daigaku Zasshi 71(6):360–370.

37. Chang Y-F (2008) Decrease of Entropy and Chemical Reactions. ArXiv08070256 Phys. Available at: http://arxiv.org/abs/0807.0256 [Accessed December 19, 2018].

38. Venegas-Gomez A (2014) The Thermodynamics of the living organisms: entropy production in the cell. ArXiv14108820 Phys. Available at: http://arxiv.org/abs/1410.8820 [Accessed December 19, 2018].

39. Humphrey T (1968) The development of the human amygdala during early embryonic life. J Comp Neurol 132(1):135–165.

40. LeDoux J (2007) The amygdala. Curr Biol CB 17(20):R868–874.

41. Gilles FH (2013) Amygdala. The Developing Human Brain: Growth and Epidemiologic Neuropathology (Butterworth-Heinemann), pp 77–79.

42. White JJ, Sillitoe RV (2013) Development of the cerebellum: from gene expression patterns to circuit maps. Wiley Interdiscip Rev Dev Biol 2(1):149–164.

43. Hadders-Algra M (2018) Early human brain development: Starring the subplate. Neurosci Biobehav Rev 92:276–290.

44. Tiemeier H, et al. (2010) Cerebellum development during childhood and adolescence: a longitudinal morphometric MRI study. NeuroImage 49(1):63–70.

45. Bernard JA, Leopold DR, Calhoun VD, Mittal VA (2015) Regional cerebellar volume and cognitive function from adolescence to late middle age. Hum Brain Mapp 36(3):1102–1120.

46. Johnson SB, Blum RW, Giedd JN (2009) Adolescent maturity and the brain: the promise and pitfalls of neuroscience research in adolescent health policy. J Adolesc Health Off Publ Soc Adolesc Med 45(3):216–221.

